# Algorithms to reconstruct past indels: the deletion-only parsimony problem

**DOI:** 10.1101/2024.10.24.620030

**Authors:** Jordan Moutet, Eric Rivals, Fabio Pardi

## Abstract

Ancestral sequence reconstruction is an important task in bioinformatics, with applications ranging from protein engineering to the study of genome evolution. When sequences can only undergo substitutions, optimal reconstructions can be efficiently computed using well-known algorithms. However, accounting for indels in ancestral reconstructions is much harder. First, for biologically-relevant problem formulations, no polynomial-time exact algorithms are available. Second, multiple reconstructions are often equally parsimonious or likely, making it crucial to correctly display uncertainty in the results.

Here, we consider a parsimony approach where any indel event has the same cost, irrespective of its size or the branch where it occurs. We thoroughly examine the case where only deletions are allowed, while addressing the aforementioned limitations. First, we describe an exact algorithm to obtain all the optimal solutions. The algorithm runs in polynomial time if only one solution is sought. Second, we show that all possible optimal reconstructions for a fixed node can be represented using a graph computable in polynomial time. While previous studies have proposed graph-based representations of ancestral reconstructions, this result is the first to offer a solid mathematical justification for this approach. Finally we discuss the relevance of the deletion-only case for the general case.

**Author summary:** An exciting frontier in evolutionary biology is the ability to reconstruct DNA or protein sequences from species that lived in the distant past. By analyzing sequences from present-day species, we aim to infer the sequences of their common ancestors —a process known as ancestral sequence reconstruction. This task has far-reaching applications, such as resurrecting ancient proteins and studying the biology of extinct organisms. However, a significant challenge remains: the lack of well-established methods for inferring past deletions and insertions —–mutations that remove or add segments of genetic code. In this paper, we present algorithms that lay the groundwork for addressing this gap. We show that finding the reconstructions involving only deletion events, while minimizing their number, can be done efficiently. Additionally, we show that all optimal solutions can be represented using specialized graphs. While previous studies have proposed graph-based representations of ancestral reconstructions, we are the first to provide a rigorous mathematical foundation for the use of these graphs.

## 1 Introduction

Biological sequences undergo mutations during evolution, and the most common ones are substitutions (when a nucleotide or amino acid is replaced by another), insertions (when a sequence is added inside another larger sequence), and deletions (when a part of a sequence is lost). Ancestral Sequence Reconstruction (ASR) is the task that consists in recovering the evolutionary history of a set of sequences related by a known phylogeny, that is the mutation events that explain the differences between these sequences. Finding where exactly these events occurred in the phylogeny amounts to determining the sequences that were present at the internal nodes of the phylogeny, that is, the *ancestral* sequences. ASR is a key task in bioinformatics, with wide-ranging applications, from protein engineering in biomedicine [1] and biotechnology [2], to protein resurrection for vaccine development [3] and paleontology [4], to the reconstruction of ancient genomes [5]. Additionally, ASR lies at the foundation of other tasks in bioinformatics, for example sequence alignment [6] and phylogenetic placement [7, 8], meaning that any improvements to the methodology for ASR are likely to benefit these other subjects.

Polynomial-time algorithms to infer past substitutions were introduced decades ago, and now form a solid basis for ASR [9, 10, 11, 12, 13, 14]. On the other hand, the inference of indels (insertions and deletions), despite multiple methodological attempts, does not have an established theoretical basis [6]. The lack of broadly adopted solutions for indels compared to substitutions can be explained by multiple factors. First, no exact algorithms that are polynomial both in the number of sequences and their length are known to solve the most biologically-relevant formulations of this problem. Thus, some algorithms are exact but have worst-case running times exponential either in the number of sequences [15, 16, 17] or their length [18, 19]. Usually, the available polynomial-time algorithms are heuristic [5, 16, 20], or they are based on simplified modeling approaches such as indel coding and its variants [21, 22, 23] or the assumption of site independence [24, 25]. Another challenge for the inference of past indels is that multiple reconstructions are often equally parsimonious/likely, meaning that it can be misleading to just return a single optimal solution. Instead, it is important to provide some insight about the other reconstructions. Exploring the diversity of solutions not only carries an additional computational cost, but it also poses the problem of how to correctly display uncertainty in the results. Even though this issue is also present for the inference of past substitutions, it is much more problematic for indels, as they can overlap each other. A graph-based solution to represent multiple reconstructions for a single node in the phylogeny, showing multiple alternative starting and ending positions for each putative gap, was recently put forward by Foley et al. [23]. Despite the problems outlined above, indel inference shows a rebirth of interest, with two parsimony-based works published in the past year [25, 26].

Similarly to substitutions, indels can be inferred using two broad approaches. The conceptually simpler one is maximum parsimony. Here, a number of formulations of the problem were already proposed two decades ago [18, 27, 15]. Although similar in spirit, these approaches differred in terms of some important modeling choices (e.g. on how to count indels between two sequences, and on whether to allow multiple introductions of the same character). The formulation by Chindelevitch et al. [27] is arguably the most faithful to the biological context, and was later adapted into number of maximum-likelihood approaches for indel inference [28, 16, 29]. It leads to a problem called the Indel Parsimony Problem (IPP), which we formally define below (Def. 5). Although the IPP is NP-hard for trees of unbounded degree, its complexity remains unknown for trees with bounded degree, in particular binary trees, and under the unit-cost model, consisting of minimizing the number of indels irrespective of their size or place in the tree.

The second broad approach for indel inference is the statistical one. In practice one assumes a probabilistic model explicitly describing the evolution of indels (e.g. [30, 31, 32, 33, 34]) and then seeks the evolutionary history or histories with highest probability [19, 16, 29, 35]. This idea can be declined either as an optimization problem or in a Bayesian context [36, 24]. For many tasks in phylogenetics, maximum parsimony has a computational advantage over statistical methods, which may rapidly become considerable as the number of sequences grows [37, 38]. It is reasonable to expect a similar advantage for indel inference. Moreover statistical approaches may benefit from the insights gained from studying the algorithmics of maximum parsimony (see, e.g., the similarities between Sankoff’s and Felsenstein’s algorithms [10, 39]).

In this paper, we focus on the specific case of parsimony-based ASR where solutions are constrained to only involve deletions. This problem, which we call the *deletion-only parsimony problem* (DPP), was first introduced by Chindelevitch et al. [27]. As we discuss more extensively in Section 7, the DPP is relevant to the solution of the general problem (the IPP mentioned above), as parts of the IPP can be optimally solved alongside this problem. The first contribution of this paper is the proposal of an exact algorithm to find all optimal solutions of the DPP. Since the number of solutions may grow exponentially in the size of the input alignment, our algorithm will also run in exponential time. However, it is easy to modify the algorithm so that it can find a single optimal solution in polynomial time. Naturally, we formally prove the correctness of this algorithm (i.e. that all, and only, the optimal solutions of the DPP are found). Secondly, we propose a polynomial alternative for the exponential-time part of the algorithm, constructing a graph for each internal node of the phylogenetic tree. Each of these graphs represents exactly the set of all reconstructions that can be found at the specified node in an optimal solution for the whole tree. These graphs are similar to the ones constructed by Foley et al. [23] and Tule et al. [26], also known as *partial order graphs* (POGs; [40]). We discuss the relation with these and other recent works in Section 7.

After this introduction, Section 2 settles the notations and definitions that are necessary to introduce the IPP and the DPP. Then, Section 3 describes our algorithm, and illustrates it step by step with a recurring example. Section 4 presents the theoretical results about the algorithm’s correctness. Then, in Section 5, we describe and illustrate our graph-based representation of optimal solutions, and the algorithm to construct it. Afterwards, in Section 6, we analyze the asymptotic complexity of all the provided algorithms. Finally, Section 7 concludes with some remarks about our original contributions, their practical relevance, and the questions left open. All the proofs of the statements in this article are given as Supporting Information, alongside some intermediate results.

## 2 Preliminaries

### 2.1 Basic notation

#### Notation for phylogenetic trees

We define a *(phylogenetic) tree T* as a binary rooted tree, whose root, set of edges, set of nodes and set of leaves are denoted by *r*(*T*), *E*(*T*),*V* (*T*) and *L*(*T*), respectively. Given two nodes *u, v* ∈ *V* (*T*), we say that *v* is a *descendant* of *u* and that *u* is an *ancestor* of *v* if the path from *r*(*T*) to *v* traverses *u*; in particular, if *v* ≠ *u* we say that *v* is a *strict descendant* of *u*, and that *u* is a *strict ancestor* of *v. T*_*u*_ denotes the subtree of *T* induced by the set of descendants of *u*. Finally, if (*p, w*) ∈ *E*(*T*) and *w* is a descendant of *p*, we say that *p* is the *parent* of *w*, and that *w* is a *child* of *p*.

#### Notation and assumptions for alignments

Given a set *X*, a (*multiple sequence*) *alignment A for X* is a table whose columns are labeled by a set of consecutive integers, called *sites*, and whose rows are in a one-to-one correspondence with the elements of *X*, called *taxa*. In particular, given *x* ∈ *X* and site *i*:

- *A*_*x*_ denotes the row of *A* corresponding to *x*;
- *A*[*i*] denotes the column *i* of *A*;
- *A*_*x*_[*i*] denotes the element of *A* at column *i* and at the row corresponding to *x*.

Although in general *A*_*x*_[*i*] may be an element of any fixed alphabet, throughout this paper we assume *A* has been preprocessed so as to be binary, i.e. *A*_*x*_[*i*] ∈ {0, 1}, where 1 and 0 are used to represent presence and absence of a character at site *i*, respectively. For any row *A*_*x*_ of an alignment, 𝟙 (*A*_*x*_) denotes the set of sites that are set to 1 (i.e. filled), that is 𝟙(*A*_*x*_) = {*k* : *A*_*x*_[*k*] = 1}.

For any alignment in input, we also assume that it has been preprocessed by adding a first column *A*[0] and a last column *A*[*m* + 1], with both *A*[0] and *A*[*m* + 1] only containing 1s, and where *m* is the number of columns in the original alignment. Finally we assume that no column *A*[*i*] contains only 0s.

#### Notation about intervals and gaps

Given two integers *i* and *j*, the set of all integers *k* with *i* ≤ *k* ≤ *j* is denoted by [*i, j*] and is called an *interval* ; Two intervals [*i, j*] and [*i*^*′*^, *j*^*′*^] are said to *overlap* if [*i, j*] ∩ [*i*^*′*^, *j*^*′*^] ≠ Ø; in particular, if [*i, j*] ⊈ [*i*^*′*^, *j*^*′*^] and [*i, j*] ⊉ [*i*^*′*^, *j*^*′*^], then [*i, j*] and [*i*^*′*^, *j*^*′*^] are said to *partially overlap*. Moreover, *A*_*x*_[*i, j*] denotes the sequence of characters at row *x* in alignment *A* between (and including) sites *i* and *j*.

##### Definition 1

(*gap*). Let [*i, j*] ⊆ [1, *m*]. *A*_*x*_[*i, j*] is a gap if all characters in *A*_*x*_[*i, j*] are 0s and *A*_*x*_[*i* — 1] = *A*_*x*_[*j* + 1] = 1.

### 2.2 Alignment extensions and indels

Alignments encountered in practice usually only contain information for the leaves of a phylogenetic tree and are thus alignments for *L*(*T*) where *T* is an unknown but estimable tree. Given a putative tree *T* and an alignment for *L*(*T*), the goal of ancestral indel reconstruction is to recover the pattern of gaps at the internal nodes of *T*. This idea underlies the following definitions.

#### Definition 2

(*alignment extension*). Let *T* be a tree and *A* an alignment for *L*(*T*). An alignment extension of *A* for *T* is an alignment *A*^+^ for *V* (*T*), such that 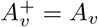 for any *v* ∈ *L*(*T*).

#### Definition 3

(*phylogenetic correctness*). An alignment extension *A*^+^ for *T* is said to be phylogenetically correct if for any two nodes *u, v* ∈*V* (*T*) and any site *i*, 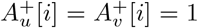 we have 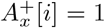 for each node *x* in the path between *u* and *v*.

The biological intuition behind phylogenetic correctness is the following. If two nodes *u, v* satisfying 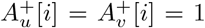 were separated by a node *x* with 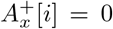, that would imply that either the character at site *i* has been introduced by two separate insertion events, or that it has first been lost with a deletion and then re-introduced with an insertion. Both of these scenarios contradict the phylogenetic interpretation of alignments, in which two characters should only be placed at the same site in an alignment if they derived from the same ancestral character. For these reasons, the problems that we consider in this paper (Definitions 5 and below) constrain alignment extensions to be phylogenetically correct. This constraint is equivalent to the common interpretation of “Dollo’s law” in phylogenetics, which asserts that no (complex) character can be introduced more than once during the course of evolution [41]. Recent works impose phylogenetic correctness by presupposing Dollo’s law [25].

The last missing ingredient before we can introduce the problems to solve is a definition of insertions and deletions, and how to count them. In a parsimony context, this can be done as in Definition 4, which is equivalent to the one by Chindelevitch et al. [27].

#### Definition 4

(*insertion, deletion, unit-cost distance*). Let *T* be a tree and (*u, v*) ∈ *E*(*T*). Let *A* be an alignment for *X* ⊇ {*u, v*}. We say that *A* has a *deletion* from *u* to *v* at interval [*i, j*] if the following statements are all verified

1. (*A*_*u*_[*i*], *A*_*v*_[*i*]) = (1, 0)
2. (*A*_*u*_[*j*], *A*_*v*_[*j*]) = (1, 0)
3. (*A*_*u*_[*k*], *A*_*v*_[*k*]) ≠ (1, 1), ∀*k* ∈ [*i, j*]
4. [*i, j*] is maximal among all intervals satisfying the three points above.

Similarly, we say that *A* has an *insertion* from *u* to *v* at interval [*i, j*] if there is a deletion from *v* to *u* at [*i, j*]. Together, insertions and deletions are called *indels*. Sites *i* and *j* are called the *endpoints* of the indel. The number of indels from *u* to *v* is also called the *unit-cost distance* between *u* and *v* and here will be noted as *d*(*A*_*u*_, *A*_*v*_).

#### Example 1.

Let *A*_*u*_ and *A*_*v*_ be defined as follows:

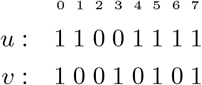

Here we have *d*(*A*_*u*_, *A*_*v*_) = 3, and more specifically a deletion at [1, 4], an insertion at {3} = [3, 3], and a deletion at {6} = [6, 6]. We use this example to illustrate the biological appropriateness of Definition 4, supported by a few informal arguments.

First, an interesting feature is that insertions and deletions can occur at overlapping intervals (e.g. [1, 4] and [3, 3] here). Indeed, other ways to define and count indels may result in considering the character losses at sites 1 and 4 as two separate deletions, implying a number of indels larger than 3 in this example [29, 15, 17]. However, this seems unjustified in a maximum parsimony context: we can imagine that at an intermediate node *x* (subdividing edge (*u, v*) into (*u, x*) and (*x, v*)) we may have *A*_*x*_ = 1 0 0 0 0 1 0 1, which can be explained with a single deletion acting on both sites 1 and 4 simultaneously from *u* to *x* (this is possible because sites 1 and 4 contain consecutive characters in the sequence at *u*). Additionally *A*_*x*_ implies a deletion at site 6 from *u* to *x* and an insertion from *x* to *v* at site 3. The existence of a scenario employing only 3 indels implies that, under maximum parsimony, *d*(*A*_*u*_, *A*_*v*_) ≤ 3.

Moreover, any definition resulting in a number of indels smaller than 3 is also biologically implausible: since from *u* to *v* some characters are lost (at sites 1, 4, 6) and some other characters are gained (site 3) there must be at least one deletion and one insertion. Let us first assume that this is possible and that the insertion occurs *before* the deletion. Then the single deletion would have to simultaneously erase sites 1, 4, 6 without erasing intermediate sites 3 and 5. This is biologically unrealistic, as a single deletion should only be allowed to erase a set of consecutive characters. Let us now assume the opposite scenario, where the insertion occurs *after* the deletion. Since the deletion must remove the characters at sites 1 and 6 and all characters that are present in *u* between them, this would result in the loss of the character at site 5, followed by its reintroduction. As argued for phylogenetic correctness, this should not be allowed to happen under a biologically-relevant modelisation of indels (see paragraph following Def. 3).

### 2.3 Problem definition

Chindelevitch et al. [27] introduced the following problem and showed that it is NP-complete on trees of unbounded degree.

#### Definition 5

(indel parsimony problem).

*Input:* A tree *T* and an alignment *A* for *L*(*T*).

*Output: A*^+^, a phylogenetically correct extension of *A*, minimizing the cost

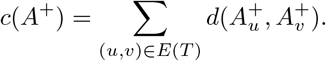

Here, we focus on a variant of this problem, where insertions are not allowed. We do this by specifying an additional constraint on *A*^+^:

#### Definition 6

(deletion-only parsimony problem – DPP).

*Input:* A tree *T* and an alignment *A* for *L*(*T*).

*Output: A*^+^, a phylogenetically correct extension of *A*, minimizing *c*(*A*^+^) under the constraint that, for all (*p, w*) ∈ *E*(*T*) where *p* is the parent of *w*, all indels from *p* to *w* must be deletions.

We will refer to the extensions *A*^+^ that satisfy the constraints of the DPP (phylogenetic correctness and all indels being deletions) as the *candidate solutions* of the DPP. Fig. 1 and 2 show an example of the input and of a candidate solution of the DPP, respectively. We note that the candidate solution in Fig. 2 is not optimal for the DPP, as we will show that there exist several solutions with a strictly lower *c*(*A*^+^).

**Figure 1:**
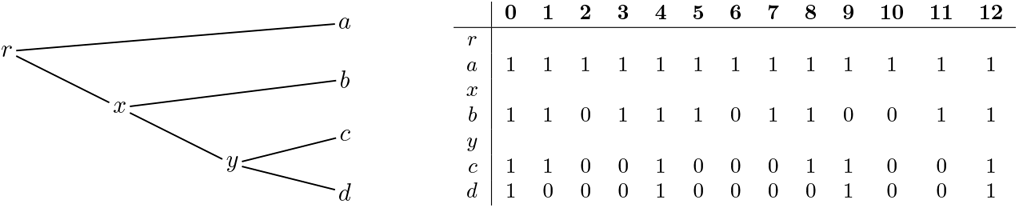
Example input of the DPP: a phylogenetic tree *T* and an alignment *A* for *L*(*T*).

**Figure 2:**
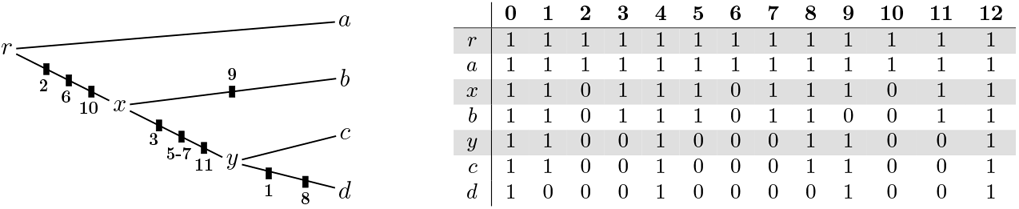
(Right:) A candidate solution *A*^+^ for the example in Fig. 1. The rows shaded in grey are those that have been added when going from the input *A* to the extension *A*^+^. They correspond to the internal nodes of *T*. (Left:) The *c*(*A*^+^) = 9 deletions in *A*^+^ are here depicted as black boxes appearing on the edges where the deletions occur, along with their endpoints. The relative order of these boxes on their edge is uninformative.

#### Observation 1.

Let *A*^+^ be a candidate solution of the DPP. Then, for every node *w* and site *k*:

- If 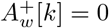 then 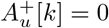, ∀*u* ∈ *V* (*T*_*w*_).
- If 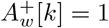 then 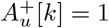, ∀*u* in the path between *r*(*T*) and *w*.
- 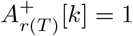.

It is easy to see that while points 1 and 2 in Observation 1 are due to the requirement that no insertions are allowed, point 3 depends on the assumption that no column of the input alignment *A* can only consist of gaps (0s). As we mentioned before, we will make this assumption throughout this paper.

## 3 The *deletion-only* algorithm

We now describe an algorithm that solves the DPP (Def. 6). The algorithm is designed to find not just one optimal solution, but all of them.

### High-level description

Our algorithm for the DPP consist of two phases. The first phase is a bottom-up procedure, starting from the leaves and proceeding up to the root of the tree, conveying information about the presence of gaps in the descendants of each node. By doing so, we can determine some gaps that are necessarily present in every optimal solution (called 0-gaps, see next paragraph), and take note of the parts that can only be determined later on. The second phase is a top-down procedure, from the root to the leaves. It allows us to resolve the parts undetermined during the first phase, sometimes with multiple choices, providing a way to generate every possible optimal solution. Algorithm 1 provides the pseudo code for our algorithm, where procedures BottomUp and TopDown are defined further below (Algorithms 3 respectively).

### Algorithm-specific notation

For each node we store a set of *labeled gaps* defined as intervals [*i, j*] ⊆ [1, *m*] labeled with one of three symbols: 0, C or P. If a gap is labeled by 0, C, P we call it *0-gap, C-gap* or *P-gap*, respectively. We explain the choice of this naming later below. If a 0-gap [*i, j*] is stored for node *u*, we also say that *u* has a 0-gap [*i, j*] or that [*i, j*] is a 0-gap at *u*. The same applies to C-gaps and P-gaps.

Remark that labeled gaps are those stored by the algorithm during its execution, and should not be confused with the gaps *tout-court* which are those that can be observed in an alignment, such as the input *A* or the output *A*^+^ (Definition 1).

### Bottom-up phase

We now describe how labeled gaps are created by the algorithm for every node *w*, in a bottom-up fashion. Algorithm 2 provides a recursive implementation of the bottom-up phase. If *w* is a leaf, its labeled gaps are set to the maximal intervals of sites filled with 0s in *A*_*w*_ and labeled as 0-gaps. In the example of Fig. 1 leaf *d* has three 0-gaps: [1, 3], [5, 8] and [10, 11].

Now let *w* be an internal node and *u* and *v* its children. For every pair of overlapping labeled gaps [*i*_*u*_, *j*_*u*_], [*i*_*v*_, *j*_*v*_] at *u* and *v*, respectively, we store the labeled gap [*i, j*] = [*i*_*u*_, *j*_*u*_] ∩ [*i*_*v*_, *j*_*v*_] for node *w* and set its label with one of the following rules.

**R1.1** If [*i*_*u*_, *j*_*u*_] = [*i*_*v*_, *j*_*v*_] and neither of them is a P-gap at *u* and *v*, then [*i, j*] is labeled as a 0-gap at *w*.

**R1.2** If [*i*_*u*_, *j*_*u*_] = [*i*_*v*_, *j*_*v*_] and only one of them is a P-gap at *u* or *v*, then [*i, j*] is labeled as a C-gap at *w*.

**R1.3** If [*i*_*u*_, *j*_*u*_] = [*i*_*v*_, *j*_*v*_] and both are P-gaps at *u* and *v*, then [*i, j*] is labeled as a P-gap at *w*.

#### Algorithm 1: DeletionOnly(*T, A*)

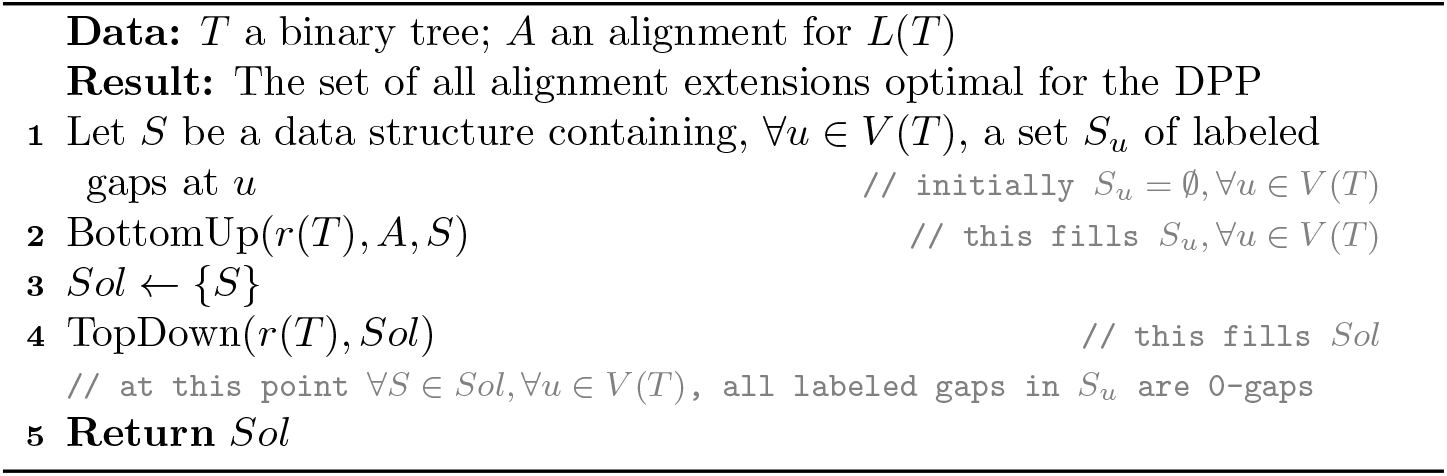

#### Algorithm 2: BottomUp(*w, A, S*)

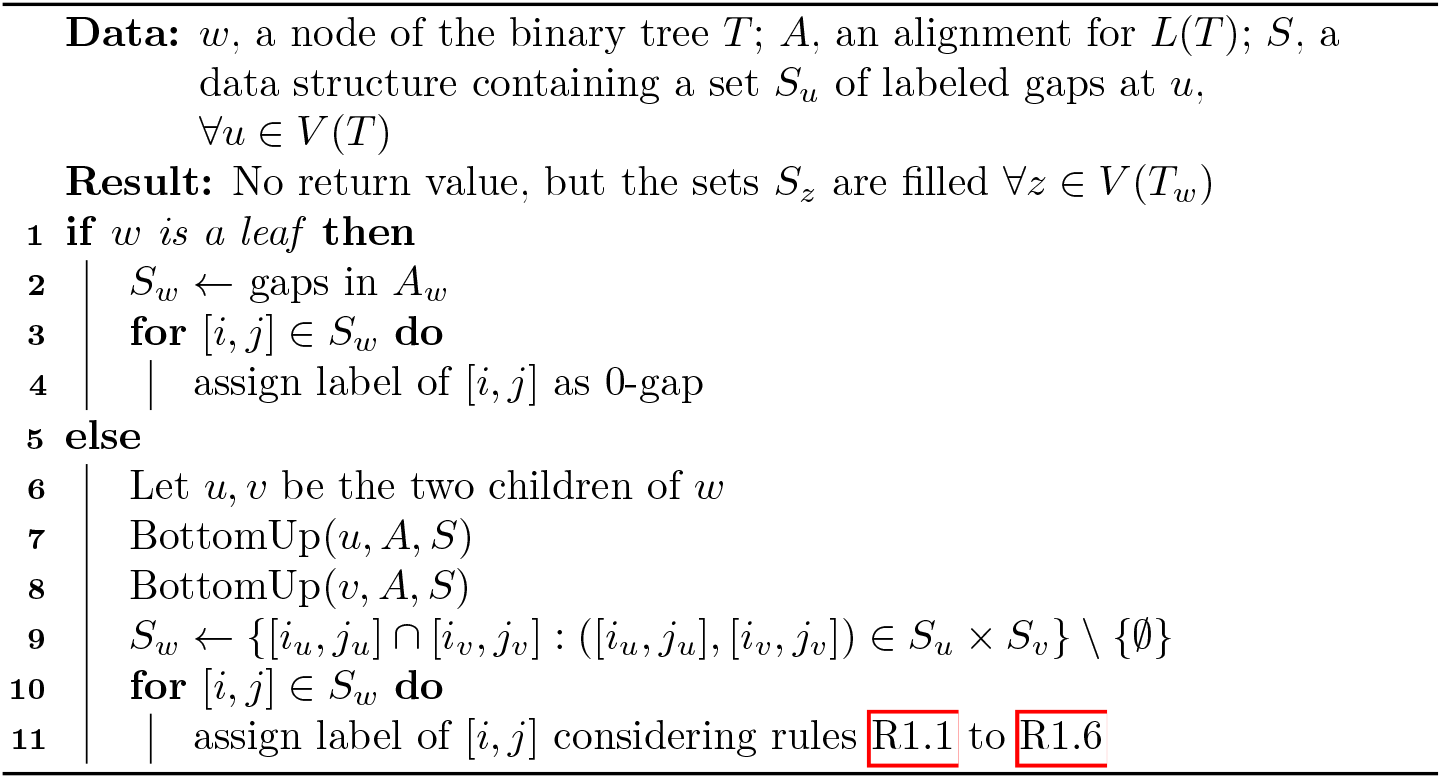

**R1.4** If [*i*_*u*_, *j*_*u*_] t [*i*_*v*_, *j*_*v*_] and [*i*_*u*_, *j*_*u*_] is not a P-gap at *u*, then [*i, j*] is labeled as a C-gap at *w*. The same applies if the roles of *u* and *v* are reversed.

**R1.5** If [*i*_*u*_, *j*_*u*_] t [*i*_*v*_, *j*_*v*_] and [*i*_*u*_, *j*_*u*_] is a P-gap at *u*, then [*i, j*] is labeled as a P-gap at *w*. The same applies if the roles of *u* and *v* are reversed.

**R1.6** If [*i*_*u*_, *j*_*u*_] and [*i*_*v*_, *j*_*v*_] partially overlap, then [*i, j*] is labeled as a P-gap at *w*.

These rules cover every case and are exclusive. In fact, R1.1, R1.2, R1.3 obviously cover all equality cases, R1.4 and R1.5 cover the strict inclusion cases, and finally R1.6 covers the partial overlap case. Table 1 displays these labeling rules as a function of the labels of [*i*_*u*_, *j*_*u*_], [*i*_*v*_, *j*_*v*_], and of whether the two gaps are identical, strictly included in one another, or partially overlapping.

**Table 1:** Depiction of how the label of [*i, j*] = [*i*_*u*_, *j*_*u*_] ∩ [*i*_*v*_, *j*_*v*_] is assigned as a function of whether [*i*_*u*_, *j*_*u*_], [*i*_*v*_, *j*_*v*_] are identical (row =), nested (rows ⫋, ⫌), or partially overlapping (row “else”), and as a function of the labels of [*i*_*u*_, *j*_*u*_] (*u* columns) and of [*i*_*v*_, *j*_*v*_] (*v* columns).

| | $u$ | $v$ | $u$ | $v$ | $u$ | $v$ |
| --- | --- | --- | --- | --- | --- | --- |
|  | 0,C | 0,C | 0,C | P | P | P |
| = | 0 |  | C |  | P |  |
| $\subsetneq$ | C | | C | | P | |
| $\supsetneq$ | C | | P | | P | |
| else | P |  | P |  | P |  |

The rationale for these rules is to guarantee the following properties. If [*i, j*] is a 0-gap at *w*, it means that *w* is either a leaf with this gap, or if we let *u, v* be the children of *w*, that there exists at least one leaf in *T*_*u*_ and at least one leaf in *T*_*v*_ such that [*i, j*] is a gap at those leaves. If [*i, j*] is a C-gap, it means that only one (but not the other) of *T*_*u*_ and *T*_*v*_ has some leaf with the same gap. Finally, if [*i, j*] is a P-gap at *w*, it means that none of the leaves that descend from *w* have this same gap. See Lemma 3 in for a proof.

Labeled gaps also have implications about an optimal solution *A*^+^ of the DPP, as shown by a few propositions stated in the next section: Proposition 1 shows that if a site *k* does not belong to any labeled gap at *w*, then we must have 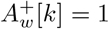. Propositio must be a gap; Proposition shows that if [*i, j*] is a 0-gap at *w*, then *k* does not belong to any labeled gap at *w*, then we must have 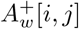 must be a gap; Proposition 3 shows that if [*i, j*] is a P-gap at *w*, then 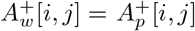 where *p* is *w*’s parent (hence the use of letter P for “parent”); finally Proposition shows that if [*i, j*] is a C-gap at *w*, then either 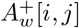 must be a gap, or 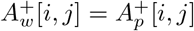 (hence letter C for “choice”).

### Example for the bottom-up phase

At the end of the bottom-up phase, the example shown in Fig. 1 produces the following labeled gaps at the internal nodes of the input tree.

Node *y* has two C-gaps, [2, 3] and [5, 7], both obtained by application of rule R1.4 on two 0-gaps at *c* and *d* (the children of *y*) with one 0-gap strictly contained in the other. Node *y* also has a 0-gap [10, 11] obtained by application of rule R1.1 on two identical 0-gaps at *c* and *d*.

Node *x* has two C-gaps, [2, 2] and [6, 6], both obtained by application of rule R1.4 on a 0-gap at *b* strictly contained in a C-gap at *y*. Node *x* also has a P-gap [10, 10] obtained by application of rule R1.6 on two partially overlapping 0-gaps at *b* and *y*.

Finally, *r* has no labeled gap, as no two labeled gaps at *a* and *x* overlap. The labeled gaps listed above are depicted in Fig. 3, although we remark that the algorithm does not need to store labeled gaps in table form.

**Figure 3:**
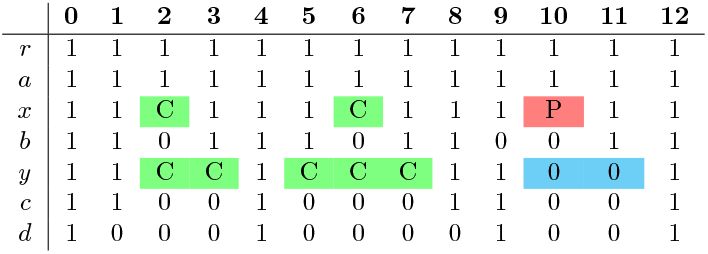
A depiction of the labeled gaps constructed by the bottom-up phase on the input shown in Fig. 1. Note that Algorithm 2 does not store labeled gaps in this form.

### Top-down phase

The goal of the second phase is to remove C-gaps and Pgaps and replace them with one or more 0-gaps. These 0-gaps, together with those already determined during the bottom-up phase, will constitute the actual gaps in the output. As we explain below, sometimes (namely when applying rule R2.2), multiple optimal choices are possible, in which case we duplicate the current solution and carry on.

Algorithm 3 provides a recursive implementation of this phase in full detail, where intermediate solutions (those still containing C-gaps and P-gaps) are represented using a data structure *S* (e.g. a map) that allows to retrieve a set *S*_*w*_ of labeled gaps for each node *w*. We visit every node of the tree in pre-order. It is easy to see that the root *r*(*T*) cannot have any labeled gaps, because otherwise the input alignment would have one or more columns only containing 0s. (See Corollary 2 in S1.) Thus, no action is needed for *r*(*T*). For every other node *w*, we process each of its C-gaps and P-gaps using one of the rules R2.1 to R2.3 below, where *p* denotes the parent of *w*.

**R2.1** If [*i, j*] is a C-gap at *w*, and [*i, j*] is a 0-gap at *p*, then [*i, j*] is reset as a 0-gap at *w*.

**R2.2** If [*i, j*] is a C-gap at *w*, and [*i, j*] is not a 0-gap at *p*, we duplicate the current solution. In the original solution we reset [*i, j*] as a 0-gap at *w*. In the duplicate, we copy at *w* all 0-gaps that are in [*i, j*] at *p*. That is, for each 0-gap [*i*^*′*^, *j*^*′*^] ⊆ [*i, j*] at *p*, we set [*i*^*′*^, *j*^*′*^] as a 0-gap at *w*, and [*i, j*] is removed from the set of labeled gaps at *w*.

**R2.3** If [*i, j*] is a P-gap at *w*, then we copy at *w* all 0-gaps that are in [*i, j*] at *p*. That is, for each 0-gap [*i*^*′*^, *j*^*′*^] ⊆ [*i, j*] at *p*, we set [*i*^*′*^, *j*^*′*^] as a 0-gap at *w*. Moreover, [*i, j*] is removed from the set of labeled gaps at *w*.

As a result of the repeated application of these rules, in the end every stored solution is encoded as a collection of 0-gaps. To produce an alignment extension *A*^+^ it suffices to set every site within a 0-gap to 0 and every other site to 1.

We remark that every time we copy at *w* all 0-gaps that are in [*i, j*] at *w*’s parent *p*, this has the effect of setting 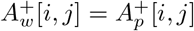. This is a consequence of the fact that no two labeled gaps [*i*_*w*_, *j*_*w*_], [*i*_*p*_, *j*_*p*_] at *w* and *p* respectively, can be partially overlapping (see Lemma 1 in S1).

#### Algorithm 3: TopDown(*w, Sol*)

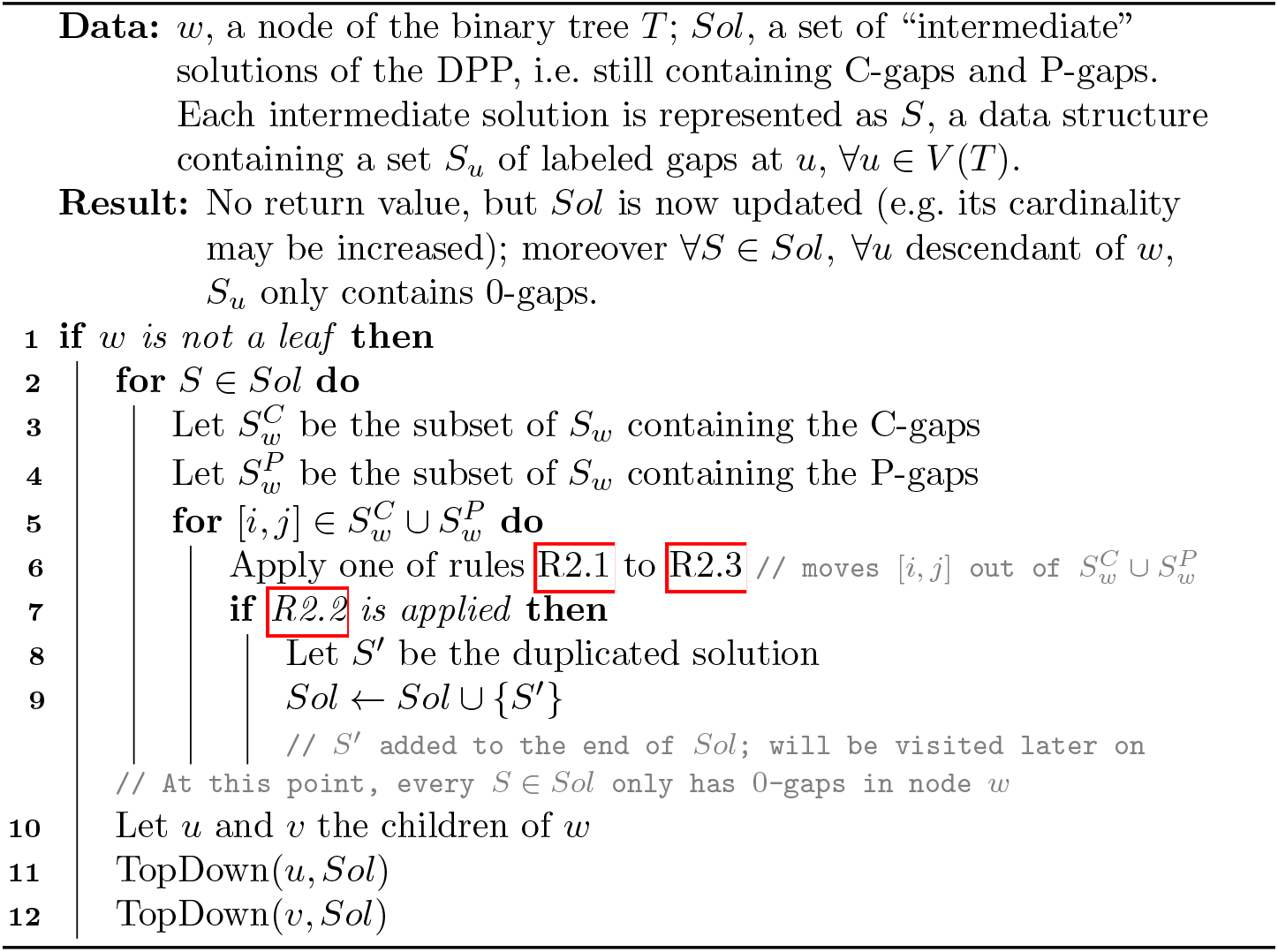

### Example for the top-down phase

The top-down phase starting from the labeled gaps shown in Fig.3 produces 16 different optimal solutions. (Because there are four C-gaps and each C-gap can be processed in two possible ways by rule R2.2.) Fig.4 shows how the top-down phase constructs one of these optimal solutions.

At *x*, both C-gaps are reset as 0-gaps by one of the two choices for rule R2.2 (steps 1,2 in Fig. 4), and the P-gap is set to 1 by rule R2.3, as *r* has no 0-gap at the interval [10, 10] (step 3). At *y*, we only apply rule R2.2: C-gap [2, 3] is replaced by 0-gap [2, 2], which is a 0-gap at *x* (in practice this sets 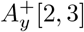 as a copy of 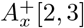 see step 4 in Fig. 4); the C-gap [5, 7] at *y* is instead reset as a 0-gap (step 5).

**Figure 4:**
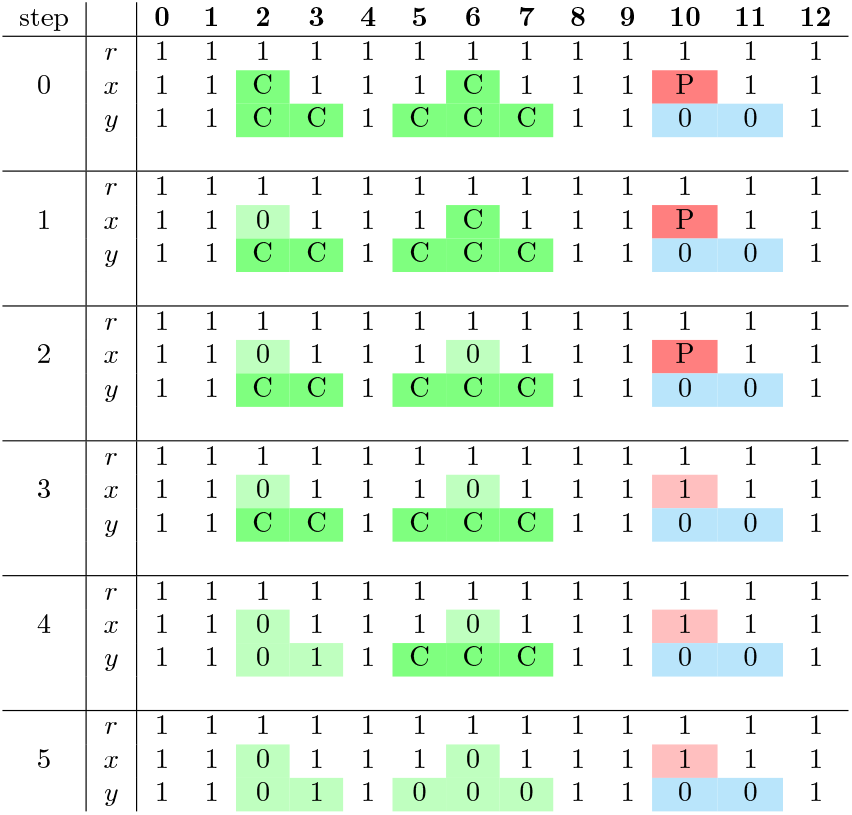
A sequence of intermediate solutions produced by the top-down phase on the input of Fig. 1, leading to an optimal solution (step 5). Each solution is represented as an alignment restricted to the internal nodes *r, x* and *y*, even though the algorithm does not store labeled gaps in this form. Colored rectangles indicate sites covered or previously covered (lighter color) by labeled gaps. At step 0, no labeled gap has been processed and the intermediate solution coincides with that produced by the bottom-up phase (Fig. 3). Each of the subsequent steps processes exactly one Cor P-gap. At steps 1, 2, 4 and 5, rule R2.2 is employed and a different choice from the one depicted could have been made.

The optimal solution produced in the end is depicted in its entirety in Fig. 5.

**Figure 5:**
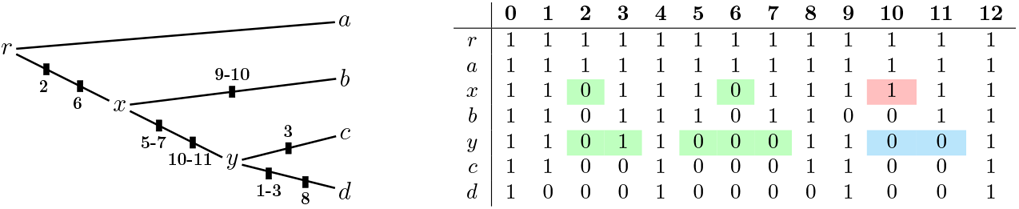
An optimal solution *A*^+^ for the example in Fig. 1, found following the steps detailed in Fig. 4. (Left:) Tree representation of *A*^+^, where each of the 8 deletions in *A*^+^ is depicted as a black box, as in Fig. 2. (Right:) Table representation of *A*^+^. Colored rectangles indicate sites previously covered by labeled gaps.

## 4. Correctness of the algorithm

In order to prove that our algorithm finds all (and only) the optimal solutions of the DPP, we need the following key results.

### Proposition 1.

At the end of the bottom-up phase, site *k* is not in a labeled gap at *w* if and only if 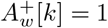 for any candidate solution *A*^+^ of the DPP.

### Proposition 2.

If [*i, j*] is set as a 0-gap at *w* during the bottom-up phase then, in every optimal solution *A*^+^ of the DPP, 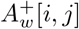 is a gap.

### Proposition 3.

If [*i, j*] is set as a P-gap at *w* in the bottom-up phase then, in every optimal solution of the DPP, 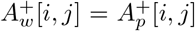 with *p* being the parent of *w*.

### Proposition 4.

If [*i, j*] is set as a C-gap at *w* in the bottom-up phase then, in every optimal solution *A*^+^ of the DPP, 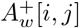 is a gap or 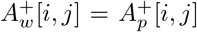. with *p* being the parent of *w*.

The four propositions above, together with a few other auxiliary results, allow us to prove the correctness of the algorithm. All proofs can be found in supplement S1.

### Theorem 1.

*A*^+^ is among the alignment extensions constructed by Algorithm 1 if and only if *A*^+^ is an optimal solution of the DPP.

## 5 Graph-based representation of solutions

Since the top-down phase can generate a number of solutions exponential in the number of C-gaps, we now describe a way to derive a compact representation of the set of all DPP-optimal reconstructions for any given node. It consists of a collection of graphs, computable in polynomial time.

For each internal node *w* ∈ *V* (*T*) \ *L*(*T*), we define a directed graph *G*_*w*_ = (*V*_*w*_, *E*_*w*_), where *V*_*w*_ = [0,*m* + 1] is the same for every *w*, and coincides with the set of sites. As for the set of arcs *E*_*w*_, it is defined by the following algorithm, which should be executed after the bottom-up phase, and replaces the top-down phase of the algorithm described in Sec. 3. We call this the *graph construction phase*.

The graph construction phase again visits the nodes of the tree with a preorder traversal. Thus, for any non-root node *w* with parent *p*, by the time *w* is traversed *E*_*p*_ has already been constructed. Moreover the set *S*_*w*_ of labeled gaps at *w* is also available.

Initially set *E*_*w*_ as follows:

**R3.0** *E*_*w*_ = {(*k, k* + 1) : *k* ∈ [0, *m*] and neither *k* nor *k* + 1 are in a labeled gap at *w*}

Then, for each labeled gap [*i, j*] at *w*:

**R3.1** If [*i, j*] is a 0-gap, then add arc (*i* — 1,*j* + 1) to *E*_*w*_.

**R3.2** If [*i, j*] is a P-gap, then for each arc (*h, k*) ∈ *E*_*p*_ with *h, k* ∈ [*i* — 1,*j* + 1], add (*h, k*) to *E*_*w*_.

**R3.3** If [*i, j*] is a C-gap, then first add arc (*i* — 1,*j* + 1) to *E*_*w*_; next, for each arc (*h, k*) ∈ *E*_*p*_ with *h, k* ∈ [*i* — 1,*j* + 1], add (*h, k*) to *E*_*w*_.

In other words, if we only consider the set of arcs in *E*_*w*_ that are between two vertices in [*i* — 1,*j* + 1], which we denote with *E*_*w*_[[*i* — 1,*j* + 1]], we have:

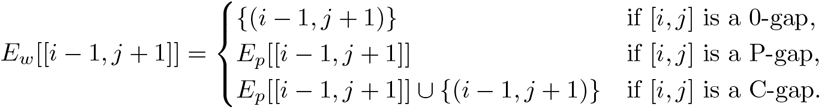

Note that since the root *r*(*T*) has no labeled gap (see Corollary 2 in S1), algorithm sets *E*_*r*(*T*)_ = {(*k, k* + 1) : *k* ∈ [0, *m*]}. the

### Correctness of the graph construction phase

Theorem 2 below states that each directed path in *G*_*w*_ from the first to the last site corresponds to one possible way of setting 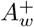 optimally, and inversely every optimal 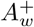 corresponds to one such path. Thus *G*_*w*_ provides a compact representation of all the optimal ancestral reconstructions for node *w*.

More formally, the graph *G*_*w*_ verifies the following property, where we recall that 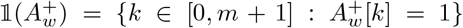. The proof can be found in supplement S2.

**Theorem 2**. Let *V* ⊆ [0,*m* + 1] and *w* ∈ *V* (*T*) \ *L*(*T*).

*V* is the vertex set of a directed path from 0 to *m* +1 in *G*_*w*_ if and only if I*A*^+^ optimal solution of the DPP such that 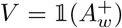.

### Example for the graph construction phase

Suppose we executed the bottom-up phase on the input of Fig. 1, producing the labeled gaps in Fig. 3. The graph construction phase starts by constructing *G*_*r*_, which (as noted above) is the trivial graph where two vertices are only adjacent if they are consecutive.

Next, *G*_*x*_ = ([0, 12], *E*_*x*_) is set to the graph shown in Fig. 6. In detail, rule R3.0 initially sets *E*_*x*_ = {(0, 1), (3, 4), (4, 5), (7, 8), (8, 9), (11, 12)} (as sites 2, 6 and 10 are in labeled gaps). Then rule R3.3 is applied to C-gap [2, 2], which leads to adding first arc (1, 3) and then arcs (1, 2), (2, 3) (copied from *G*_*r*_). Next, rule R3.3 is applied to C-gap [6, 6], which similarly leads to adding arcs (5, 7), (5, 6), (6, 7) to *E*_*x*_. Finally, rule R3.2 is applied to P-gap [10, 10], which results in adding arcs (9, 10), (10, 11) (copied from *G*_*r*_). This completes the construction of *G*_*x*_.

**Figure 6:**
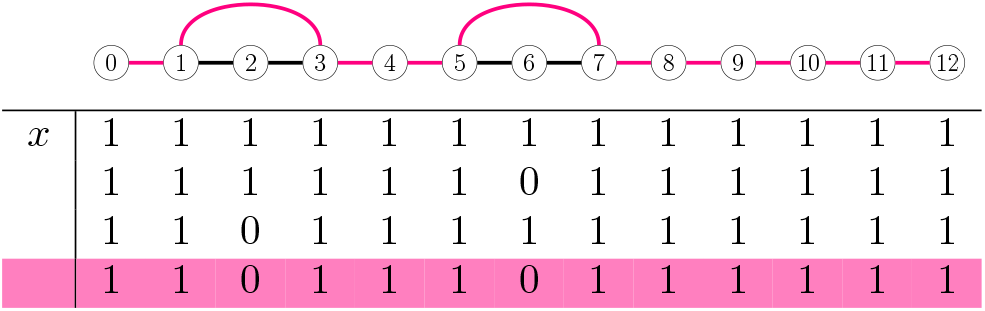
Graph representation of all possible 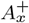, where *A*^+^ is constrained to be an optimal solution for the example in Fig. 1. (Top:) The graph *G*_*x*_ obtained by the graph construction phase, with one path highlighted. (Bottom:) The four possible 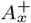 when 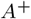 is an optimal solution. By Theorem 2, there is a one-to-one correspondence between these 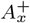 and the paths from vertex 0 to vertex 12 in *G*_*x*_. The path highlighted in *G*_*x*_ corresponds to the highlighted 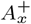.

Next, *G*_*y*_ = ([0, 12], *E*_*y*_) is set to the graph shown in Fig. 7. In detail, rule R3.0 initially sets *E*_*y*_ = {(0, 1), (8, 9)}, as the labeled gaps at *y* are [2, 3], [5, 7], [10, 11]. Then rule R3.3 is applied to C-gap [2, 3], which leads to adding to *E*_*y*_ arc (1, 4) and then all the arcs in *E*_*x*_[[1, 4]]. Next, rule R3.3 is applied to C-gap [5, 7], which leads to adding to *E*_*y*_ arc (4, 8) and then all the arcs in *E*_*x*_[[4, 8]]. Finally, rule R3.1 is applied to 0-gap [10, 11], which results in adding arc (9, 12) to *E*_*y*_. This completes the construction of *G*_*y*_, and the graph construction phase terminates.

**Figure 7:**
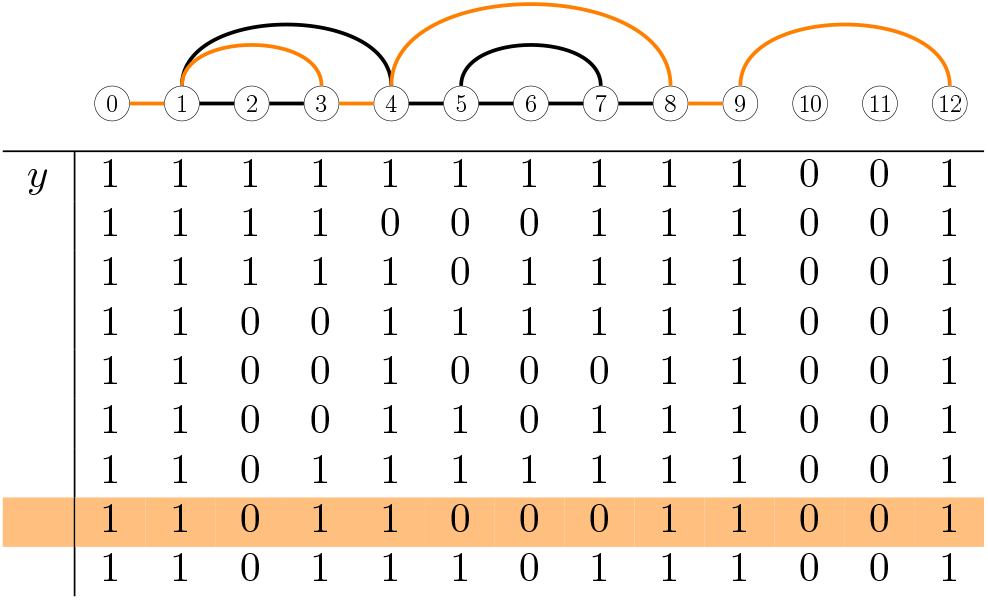
Graph representation of all possible 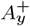, where 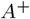 is constrained to be an optimal solution for the example in Fig. 1. (Top:) The graph *G*_*y*_ obtained by the graph construction phase, with one path highlighted. (Bottom:) The nine possible 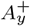 when 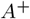 is an optimal solution. The path highlighted in *G*_*y*_ corresponds to the highlighted 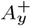.

To illustrate Theorem 2, we focus on *G*_*y*_ (Fig. 7). Theorem implies that the set of paths in *G*_*y*_ from 0 to 12 is in a one-to-one correspondence with the set of binary strings that can be obtained as 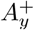 for some optimal solution *A*^+^.

To verify this, we can imagine deriving all 16 optimal solutions *A*^+^ and then taking the row 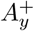 in each of them. Doing so produces 9 different values for 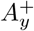, the binary rows depicted in Fig. 7. (Sometimes pairs of optimal solutions that have different 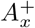 may have the same 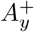, which explains why from 16 optimal solutions we only get 9 different 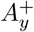.) It is easy to check that each of these 9 values for 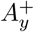, when translated as 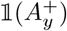, gives the vertex set of a path from 0 to 12 in *G*_*y*_. Conversely, each path in *G*_*y*_, when converted into the binary string that has a 1 at site *k* if the path traverses *k*, corresponds to one of the 9 rows shown in Fig. 7.

## 6 Complexity

Here we state the main results about the complexity of our algorithms for the DPP. Full proofs can be found in supplement S3. The asymptotic complexity will be expressed as a function of the following parameters:

- *n*, the number of leaves of *T* or equivalently the number of rows of *A*;
- *s*, the number of optimal solutions of the DPP;
- *b*, the number of boundaries in *A*, where a *boundary* is defined as a pair of consecutive sites (*k, k* + 1), such that *A*[*k*] ≠ *A*[*k* + 1], i.e., their corresponding columns differ;
- *m*, the number of columns in *A* (excluding the first and last column which we only added for convenience of notation).

For the analysis of Algorithm 1, we assume that both the input alignment *A* and the output alignment *A*^+^ are encoded in *gap form*, instead of the table form presented in the Preliminaries (sec. 2.1). The gap form of an alignment *A* consists of encoding each row of *A* as the set of gaps it contains. For example, the input alignment of Fig. 1 can be encoded as *A*_*a*_ = Ø, *A*_*b*_ = {[2, 2], [6, 6], [9, 10]}, *A*_*c*_ = {[2, 3], [5, 7], [10, 11]}, *A*_*d*_ = {[1, 3], [5, 8], [10, 11]}.

We assume that alignments are represented in gap form because, as a result, running times and memory requirements only depend on parameter *b* rather than *m*. In practice, for many alignments, we can expect *b* to be much smaller than *m*. We can now state the first of the two main results of this section.

**Theorem 3**. Assume that the input alignment *A* and every solution *A*^+^ in output is represented in gap form. Then Algorithm 1 runs in *O*(*nbs*) time.

Note that an obvious bound is *s* ≤ 2^*c*^, where *c* denotes the number of Cgaps at the end of the bottom-up phase. Although the worst-case complexity of Algorithm 1 is exponential in *c*, we can avoid the exponential factor in two ways. First, we can simply keep only one solution, which can be achieved by removing the duplication in rule R2.2 and skipping lines 7 – 9 in Algorithm 3. The complexity then becomes *O*(*nb*).

Alternatively the top-down phase can be replaced by the graph construction phase described in Sec. 5. We now analyze the complexity of this algorithm. A key observation here is given by the following proposition.

### Proposition 5.

For any internal node *w, G*_*w*_ is a planar graph.

Proposition 5 can be used to obtain the other main complexity result:

**Theorem 4**. The algorithm that first runs the bottom-up phase and then the graph construction phase runs in *O*(*nm*) time.

Note that the complexity in Theorem 4 depends on *m* rather than on *b*. This is only because, for simplicity, we have chosen to describe the graph construction phase so that the graphs have [0,*m* + 1] as vertex set. It is possible to see that it is feasible to modify the graph construction phase so that each graph has a vertex set of size *O*(*b*). Doing so would allow us to obtain an algorithm running in *O*(*nb*) time. We leave this as future work.

## 7 Discussion

### Novel contributions and related works

We have focused on a version of the Indel Parsimony Problem where only deletions are allowed (the DPP; Definition 6). We have described an exact algorithm that constructs all the optimal solutions for the DPP, and proved it is correct. While the algorithm’s running time may be exponential in the size of the input, this is only due to the exponential number of solutions. The algorithm can be easily changed to only find a single optimal solution in polynomial time (e.g. skip lines 7 – 9 in Algorithm 3). We note that an algorithm to find an optimal solution of the DPP was also proposed by Chindelevitch et al. [27]. Unfortunately, the pseudo-code provided in that paper appears to be incorrect, although the main underlying ideas are sound. See supplement S4, where we copied the pseudocode and provide a simple example where it fails to return an optimal solution. Importantly, the main difference with our work is that we seek the whole set of optimal solutions, a goal which significantly complicates the task.

Two recently published parsimony-based works also deserve some discussion [25, 26]. Iglhaut et al. [25] proposed an extension of Fitch’s algorithm [9] that handles insertions and deletions. The proposed algorithm is not intended to solve any explicit formulation of the parsimony principle for indels. Importantly, like in the original algorithm by Fitch, ancestral reconstruction is executed one site at a time, independently. Second, Tule et al. [26] present a Mixed-Integer Programming (MIP) formulation of indel inference, which provides an exact solution, albeit not polynomial in the worst case. However, their formulation is radically different from that of the Indel Parsimony Problem [27]. For example, phylogenetic correctness is not imposed, and the number of indel events between the reconstructions for two adjacent nodes can be very different (e.g. Tule et al. [26] count three separate indel events between 1 0 1 0 1 0 1 0 1 and 1 0 0 0 0 0 0 0 1 whereas Def. 4 only one).

Back to our paper, since there is no reason to favor one optimal scenario over an equally-parsimonious one, we consider it important to conceive ways to represent compactly the diversity of the reconstructed solutions. Our other main contribution is that we have shown that, in the context of the DPP, it is possible to efficiently construct a graph *G*_*w*_ representing the set of optimal reconstructions 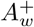 for any internal node *w* of the phylogeny. While the software GRASP does indeed construct similar graphs [23], we note that there is no proven global optimality guarantee for the reconstructions represented by the graphs of GRASP. We believe that our result (i.e. Theorem 2) is the first to provide a mathematical justification for the use of partial order graphs [40] in the context of ancestral sequence reconstruction. More recently, Tule et al. [26] also infer a graph for each internal node, but each of these graphs only has a single path from the first to the last site, meaning that it cannot be used to represent multiple reconstructions.

Finally we have derived the asymptotic complexity for all our algorithms. These algorithms are asymptotically optimal, because their running time is linear in the size of their inputs, or in the size on memory of its intended output (the set of all optimal solutions, or the graphs representing them).

### Practical relevance of the deletion-only case

Although this may not be evident at first sight, the relevance of deletion-only case extends beyond the solution of the DPP. In fact, algorithms to solve the DPP can be used to solve (parts of) many instances of the general problem (the IPP; Definition 5). This is due to three observations, which first we state informally and then illustrate with an example below.

- Many instances of the IPP can be solved with a divide-and-conquer approach, by subdividing the problem into subproblems, so that solving the original problem is equivalent to solving the subproblems and then combining the solutions for the subproblems. A way to do this was shown by Chindelevitch et al. [27], and another one is implicit in the work by Fredslund, Hein, and Scharling [18].
- The IPP is *time-reversible*, in the sense that if the input tree is rerooted on a node different from the original root, then every candidate solution retains its cost. As a consequence, optimal solutions of the IPP (excluding the reconstruction for the root) are independent of the position of the root.
- Recall that phylogenetic correctness may constrain an internal node *x* to have many of its sites equal to 1. Now suppose that there exists a node *x* such that *A*^+^ must only consist of 1s, for any candidate solution *A*^+^ of the IPP. Then if we re-root the tree in *x*, none of the indels in any candidate solution *A*^+^ can be an insertion (an insertion implies the existence of an edge (*p, w*) where *p* is the parent of *w*, such that 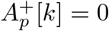 and 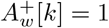 which implies 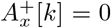). But then we can solve the IPP by solving the DPP on the re-rooted tree with one of our algorithms. The following example shows how these three ideas can be combined. Consider instance (*T, A*) of the IPP in Fig. 8, where for simplicity we do not show the dummy columns *A*[0] and *A*[5]. While at first sight there is no reason why the deletion-only case would be relevant to solve this example, it turns out that we can indeed use our algorithms for the DPP to solve it.

**Figure 8:**
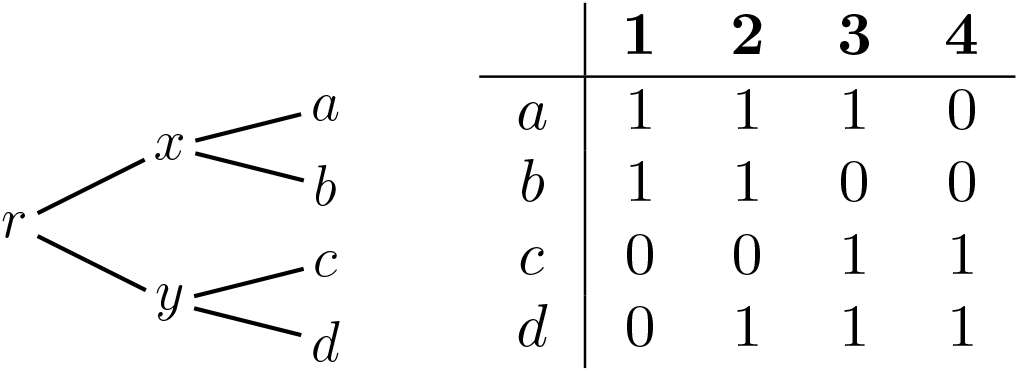
An instance (*T, A*) of the IPP that can be solved by applying the algorithms for the DPP (see the main text).

First, this is a case where we can decompose the problem in two subproblems, i.e., an example of point 1 above. Namely, we can split the input alignment in two sub-alignments *A*_1_ and *A*_2_, where *A*_1_ consists of columns 1 and 2 of the original alignment, while *A*_2_ contains columns 3 and 4. The solutions for the instance in Fig. 8 can be obtained by combining the solutions for the IPP on (*T, A*_1_) and those for the IPP on (*T, A*_2_). This can be proven by applying previous results [27, 18], but the main idea here is that no indel can occur at an interval [*i, j*] ⊇ [2, 3], for the following reasons. Phylogenetic correctness imposes 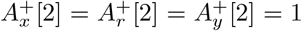 and also 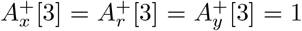 this implies that if an indel involves sites 2 and 3, then it must be a deletion from one of *x* or *y* to a leaf; but none of the leaves has two 0s at sites 2 and 3, and so we cannot have any deletion at [*i, j*] ⊇ [2, 3]. Thus, for any candidate solution, every indel must be within one of the two sub-alignments *A*_1_ or *A*_2_, which implies the decomposability of the original IPP into the subproblems (*T, A*_1_) and (*T, A*_2_).

Because (*T, A*_1_) and (*T, A*_2_) are symmetric, we can just concentrate on the instance (*T, A*_1_) of the IPP. Here phylogenetic correctness implies that every sequence on the path between *a* and *b* must have 1s at both sites 1 and 2. Thus, if we create a new node *r*^*′*^ subdividing edge (*x, a*), we know that we must have 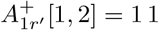. We then re-root the tree in *r*^*′*^, which gives us a new rooted tree *T* ^*′*^ shown in Fig. 9. For the reasons explained in point 2 above, solving the IPP on (*T, A*_1_) is equivalent to solving the IPP on (*T* ^*′*^, *A*_1_). (Excluding the ancestral reconstruction for the root, which we discuss below.)

**Figure 9:**
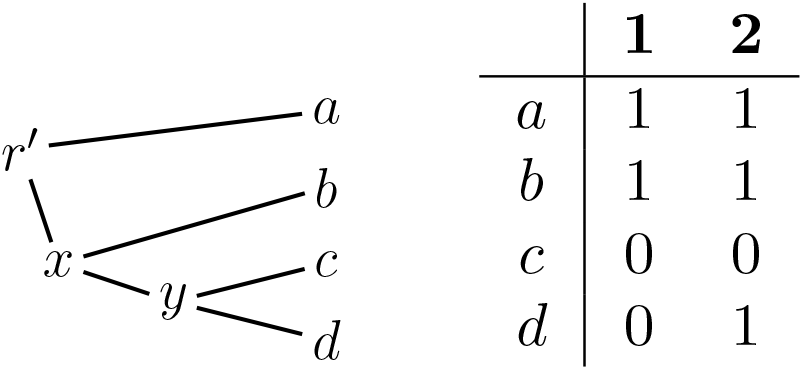
An instance (*T* ^*′*^, *A*_1_) of the DPP that provides a solution of the first half of the IPP problem in Fig. 8. This instance has two optimal solutions with 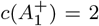. Both solutions have 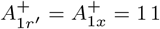, but they differ at node *y* where they have 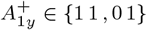.

But in turn, solving the IPP on (*T* ^*′*^, *A*_1_) is equivalent to solving the DPP on (*T* ^*′*^, *A*_1_), because of 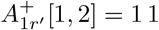 and for the reasons explained in point 3. Hence, we can use our algorithm for the DPP to solve the IPP on (*T, A*_1_).

It is now easy to check that, for the instance in Fig. 9, our algorithm gives the two solutions with 1 1 at node *x* and with one of {11, 0 1} at node *y*. By symmetry the IPP on (*T, A*_2_) has the two solutions with one of {11, 1 0} at node *x* and with 1 1 at node *y*. Combining each solution for one sub-problem with each solution for the other sub-problem gives four solutions for the original IPP on (*T, A*) (Fig. 8). They are the ones with 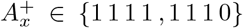 and 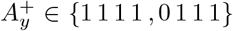. As for the root, it is easy to see that setting it equal to one of the reconstructions for its children always leads to an optimal solution. Thus, here we can set 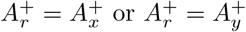.

### Open problems and future work

We note that other papers consider formulations of the IPP that are more general than ours (Definition 5). In these formulations, each indel contributes to *c*(*A*^+^) with an affine cost *α* + *βl*, where *l* is the length of the indel [18, 27, 26]. Additionally, the cost of each indel may also depend on the branch where it occurs, allowing to consider branch lengths [27]. In this paper we consider a simpler version of the IPP where each indel contributes the same unit cost (i.e. *β* = 0), irrespective of its length, or of the branch where it occurs. Note that the computational complexity result given by Chindelevitch et al. [27] implied the NP-hardness of two versions of the IPP:

- the IPP on trees with unbounded degree under the unit-cost model,
- the IPP on binary trees with branch-dependent indel costs.

This notably leaves open whether it is NP-hard to solve the IPP on binary trees and unit costs. In fact, we have chosen to tackle the DPP for binary trees and unit costs because our ideas may have an impact on this version of the IPP of unknown complexity.

While the most immediate idea for future work may be to extend our algorithms to the DPP for non-binary trees and/or affine costs, we believe that the most important avenue is to investigate whether our approach can be leveraged to also solve the general IPP on binary trees and unit costs. Alternatively, showing the NP-hardness of this problem would be an important contribution to the field.

## Notes

### Competing Interest Statement

The authors have declared no competing interest.

